# Distinct oncogenes drive distinct genome and epigenome alterations in human mammary epithelial cells

**DOI:** 10.1101/344465

**Authors:** Claire Fonti, Anne Saumet, Amanda Abi-Khalil, Béatrice Orsetti, William Jacot, Elouan Cleroux, Ambre Bender, Michael Dumas, Emeline Schmitt, Jacques Colinge, Michael Weber, Claude Sardet, Stanislas du Manoir, Charles Theillet

## Abstract

Gene expression differences, combined with distinct patterns of genomic rearrangements and epigenetic modifications, have laid the bases of molecular classification of breast cancer. Different molecular subtypes are thought to originate from different cell lineages in the mammary gland, but the early activation of an oncogene could also play a role. It is, however, difficult to discriminate the respective inputs of oncogene activation or cell type of origin in the natural history of the tumor. In this work, we have designed an experimental strategy aiming at determining whether activation of distinct oncogenic pathways in human mammary epithelial cells (HMEC) could lead to different patterns of genetic and epigenetic changes. We show that initial activation of CCNE1, WNT1 and RASv12, which activate distinct oncogenic pathways, in shp53 immortalized HMECs results in different and reproducible profiles of mRNA and miRNA expression, copy number alterations (CNA) and DNA methylation modifications. Noticeably, HMECs transformed by RAS bore very specific profiles of CNAs and DNA methylation, clearly distinct from those shown by CCNE1 and WNT1 transformed HMECs.

Genes impacted by CNAs and CpG methylation in the RAS and the CCNE1/WNT1 clusters showed clear differences, illustrating the activation of distinct pathways. Our data show that early activation of distinct oncogenic pathways leads to active adaptive events resulting in specific sets of CNAs and DNA methylation changes. We, thus, propose that activation of different oncogenes could have a role in reshaping the genetic landscape of breast cancer subtypes.

**Author summary:** Genetic and epigenetic changes are at the center of cancer development. Breast cancer molecular subtypes are defined on differences in genetic and epigenetic profiles and it is generally assumed these subtypes originate from different cell lineages in the mammary gland. We propose that founding oncogenic mutations could also have an impact. To address this question, we designed an experimental model, based on the ectopic expression of different oncogenes in human mammary epithelial cells (HMEC), and monitored genetic and DNA methylation changes occurring at different stages of cell transformation. We show that transformation of HMEC by distinct oncogenes resulted in clearly different and reproducible patterns of genetic and DNA methylation changes. Genes whose expression was modified by either CNAs or CpG methylation were consistent with the dominant pathways activated and reflected the phenotypes in the respective models. We propose that DNA methylation and CNA changes correspond to adaptive responses to the activation of the oncogenic pathways. Our data strongly suggest that early activation of distinct oncogenic insults will not only impinge on the phenotypic characteristics of the resulting tumors, but also have a strong impact on their genomic and epigenetic landscapes.

## Introduction

Genetic instability lies at the core of neoplastic development with up to 85% of human cancers showing loss of chromosome integrity at varying levels such as aberrant copy numbers and aneuploidy. Chromosomal instability has been observed in early stages of cancer [1] and rearrangement intensity correlated with disease aggressiveness [2]. In addition to structural defects, cancer genomes undergo important epigenetic changes occurring at the chromatin and DNA levels [3]. At the DNA level, cancer associated epigenetic modifications involve genome wide cytosine methylation changes corresponding to demethylation of repetitive DNA sequences and hypermethylation of CpG enriched sequences. While demethylation of repetitive DNA has been proposed to favor chromosomal instability, hypermethylation of CpG rich promoter sequences has been associated with gene expression changes[4,5].

Genetic instability results in stochastically occurring aberrations, of which a fraction will be selected according to the survival or growth advantage they confer to the cells. Hence, profiles of somatically acquired genetic and epigenetic changes and associated RNA expression patterns in tumors reflect the combined interactions of genetic instability and selective pressure. As a consequence, recurrent profiles of genomic and epigenetic aberrations should in most cancers be structured around a set of anomalies that confer maximum advantage in a given tissue and environment. Noticeably, cancers of distinct anatomical origins exhibit quite different profiles of genomic and epigenetic anomalies [6]. Recent work by Sack and coworkers [7], who screened human open reading frame libraries in 3 different cell types to identify proliferation drivers, elegantly showed the existence of a tissue specific bias in the selection of gain of function and, conversely, the counter-selection of suppressor genes.

In breast cancer, molecular subtypes were defined on the basis of RNA expression, as well as of genomic anomalies and DNA methylation differences [8-11]. Although definitive proof is still missing, it is generally proposed that the genetic and epigenetic differences in different breast tumor subtypes are dictated by distinct cell types of origin [8,9]. Early activation of distinct oncogenic pathways in a single cell type could also have an impact on genomic and epigenetic changes and induce the selection of anomalies functionally coherent with the activated pathway [12-15]. This is supported by studies showing that expression of RASv12 and of BRAFv600 resulted in the transcriptional repression and hypermethylation of distinct gene sets, involving different cascades of repressors and DNA methylases [16,17]. Yet, it has not been experimentally demonstrated that distinct oncogenic events could lead to specific genomic rearrangements.

In this work, we sought to determine the impact of the early activation of distinct oncogenic pathways on genomic and epigenetic changes in immortalized human mammary epithelial cells (HMECs). To this aim, we overexpressed by retroviral transduction three oncogenes WNT1, CCNE1 and RASv12, known to activate different oncogenic pathways, in shp53 immortalized human HMECs and monitored epigenetic and genetic changes at different steps of cancer progression. The sequence of genetic and epigenetic alterations accompanying the transition between the normal and transformed states show that activation of these distinct oncogenes leads to the emergence of distinct and specific profiles of changes. These results thus support a model in which genetic and epigenetic changes in cancer cells reflect adaptive responses to the oncogenic driver.

## Results

### Establishment of the cellular model of stepwise transformation

Primary human mammary epithelial cells (HMEC) were isolated from fresh mammary tissue obtained from donors undergoing reductive plastic surgery (details in Supplementary Methods). We obtained 3 stable cell cultures that could be propagated for at least 15 passages, before cells started showing signs of senescence such as reduced proliferation, enlarged morphology with large vacuoles and positive β-galactosidase staining (S1A Fig). Of the 3 primary HMEC lines, we selected the R2 line to establish our models of stepwise cell transformation. As depicted in Fig 1A, cells were genetically modified by sequential retroviral transductions with defined genetic elements. In the first (immortalization) step, we transduced a vector expressing an shRNA targeting the *TP53* gene (designated shp53 hereafter), which plays a key role in the senescence barrier. The shp53 efficiently knocked-down p53 protein expression and signaling (S2 Fig) and gave rise to rapidly growing cultures with a short lag period after infection. We derived three stable shp53 HMEC sublines, which we will refer to as R2*^shp53^* and used as models in this work. The three shp53 sublines showed reduced levels of β-galactosidase staining, re-expression of the endogenous *hTERT* and stabilization of telomere length, indicating that the senescence program had been overcome (Fig 1B-C, S1B-C Fig). In the second step, two of these R2*^shp53^* sublines were independently transduced with vectors expressing either the *CCNE1*, *WNT1* or *HRAS^v12^* oncogenes. These oncogenes were selected because of their known transformation potential in human cells and because they belonged to distinct signaling pathways. Oncogene expressing cells were expanded in culture and several independent sublines were derived for each oncogenic situation (R2^shp53-WNT1^, R2^shp53-CCNE1^, R2^shp53-RAS^). In the next step, each subline was seeded in soft agar to determine anchorage independent growth as a standard read out of *in vitro* transformation (S1D Fig). Cell clones were isolated from soft agar foci and expanded (R2^shp53-WNT1.SA^, R2^shp53-CCNE1.SA^, R2^shp53-RAS.SA^). Hereafter, we will refer to R2^shp53^ transduced with an oncogene as pre-transformed for cells before soft agar cloning and as transformed after soft agar cloning. In total, we established 16 HMEC sublines (S appendix) corresponding to different steps of cancer transformation; immortalized (R2^shp53^), pre-transformed (R2^shp53-oncogene)^ and transformed (R2^shp53-oncogene.SA^).

**Fig. 1:**
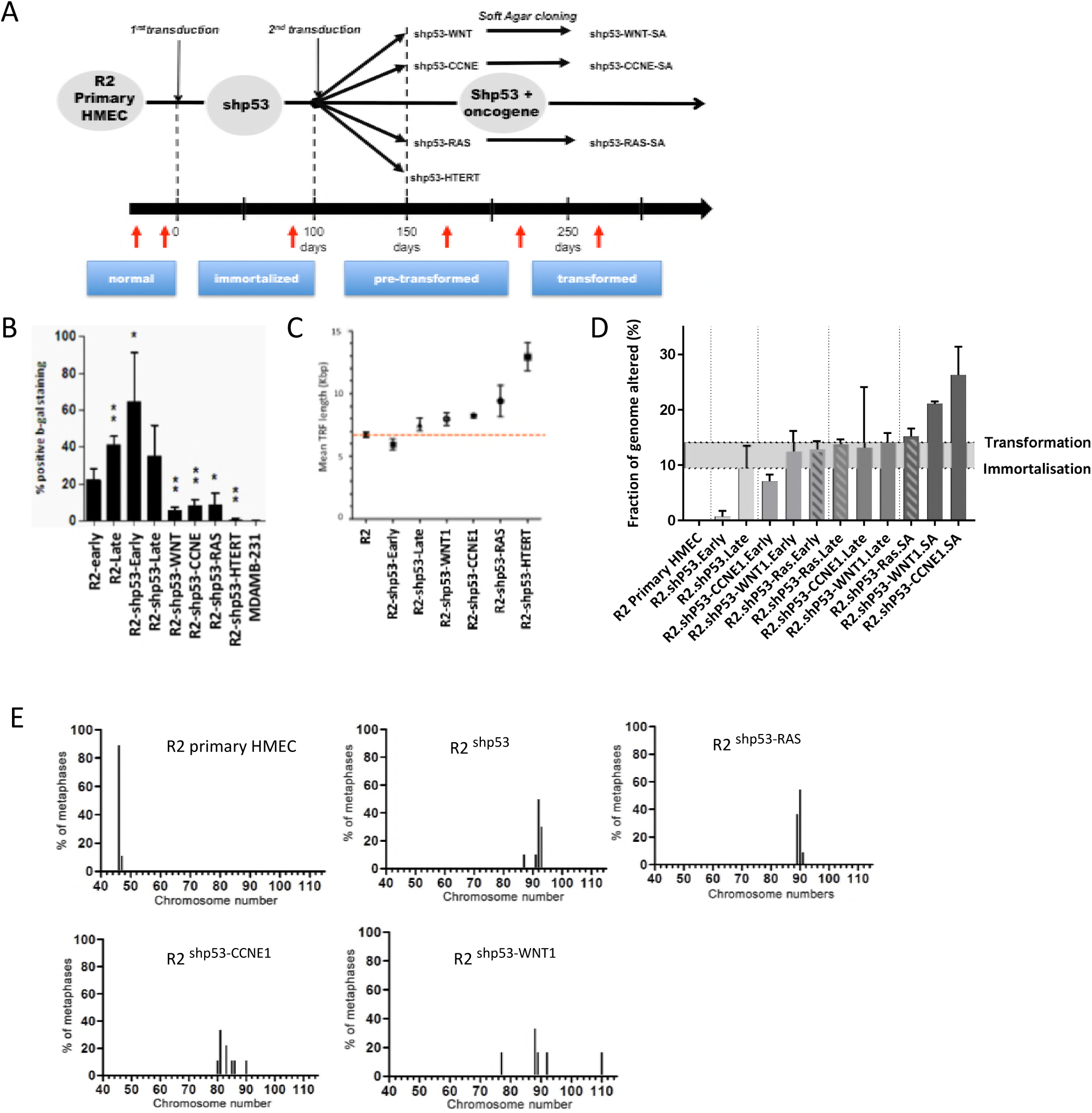
Stepwise transformation HMEC models. **A:** experimental scheme of the subline production. **B**: percentage of B-galactosidase positive cells in the different HMEC variants. **C**: mean length of telomeres estimated by Southern blotting show a stabilization of telomere length in R2 ^shp53.Late^ and transformed HMECs. **D**: fraction of the genome involved in Copy Number Alterations. Anova Multiple Comparisons test showed significant increase in genomic fraction involved in CNA between R2 primary HMEC and oncogene transduced sublines (R2 vs. R2.shP53-Ras p=0.0149; R2 vs. R2.shP53-CCNE1 and shp53-WNT1 p=0.0193). Comparison of pretransformed and transformed (after Soft Agar) sublines showed that genomic fraction involved in CNA were significantly increased in shp53-CCNE1.SA and shp53-WNT1.SA cells (shP53-CCNE1/shp53-WNT1 vs. R2.shP53-CCNE1.SA/shp53-WNT1.SA p= 0.0021) but not in shp53-RAS.SA cells (shP53-Ras vs. shP53-Ras.SA p= 0.9835). **E**: chromosome numbers were estimated on chromosome spreads. At least 50 karyotypes were scored in each subline.

### Genetic instability in the immortalized and transformed cell lines

Of the 16 established sublines, 8 were characterized at early and late passages. We first analyzed the primary, immortalized, pre-transformed and transformed HMEC sublines by array-CGH. Using the fraction of the genome involved in copy number alterations (CNAs) as a metric, it was noticeable that the level of genetic instability gradually increased between the immortalized R2^shp53^ and the transformed R2^shp53-WNT1.SA^, R2^shp53-CCNE1.SA^ and R2^shp53-RAS.SA^ sublines (Fig 2A). Indeed, the fraction of the genome involved in CNA increased from 2 to 6% in R2^shp53^ HMECs between early (50 days after transduction) and late passages (300 days). CNAs doubled (12-13% of the genome) in R2^shp53-WNT1^, R2^shp53-CCNE1^ and R2^shp53-RAS^ in comparison to R2^shp53^ HMECs and peaked in R2^shp53-WNT1.SA^ and R2^shp53-CCNE1.SA^ at 22 and 27% of the genome respectively (Fig 1D). Interestingly, the patterns of CNA progression were coherent with ploidy changes in these sublines. Indeed, chromosome counts performed on metaphase spreads revealed that R2^shp53^ HMECs presented 92 chromosomes indicative of tetraploidy. Remarkably, while all R2^shp53-RAS^ HMECs clones remained strictly tetraploid, R2^shp53-CCNE1^ and R2^shp53-WNT1^ sublines became aneuploid with chromosome counts ranging from 78 to 110 (Fig 1E). To determine whether increased CNA levels were associated to an elevation of genetic stress, we performed immunofluorescence staining of γH2Ax and 53BP1 in R2, R2^shp53^ and R2^shp53-oncogene^ HMECs (S3A-B Fig). Two types of γH2Ax staining patterns were observed; nuclear foci, considered as markers of DNA breaks, and pan-nuclear staining, which has been proposed to reveal widespread replication stress in the absence of double strand breaks [18]. In immortalized R2^shp53^ HMECs, we predominantly observed pan-nuclear γH2AX staining and low levels of γH2Ax and 53BP1 foci, suggesting an elevation of genetic stress but low levels of DNA breaks in these cells. This contrasted with R2^shp53-CCNE1^ and R2^shp53-WNT1^, which essentially displayed γH2Ax and 53BP1 nuclear foci (S3A-B Fig), consistent with a distinguishable increase of DNA breaks in these cells confirmed by comet-assay (S3C-D Fig). Of note, R2^shp53-RAS^ showed distinctly lower levels of γH2Ax and 53BP1 nuclear foci and low levels of DNA breaks by comet-assay relative to R2^shp53-CCNE1^ and R2^shp53-WNT1^.

**Fig. 2:**
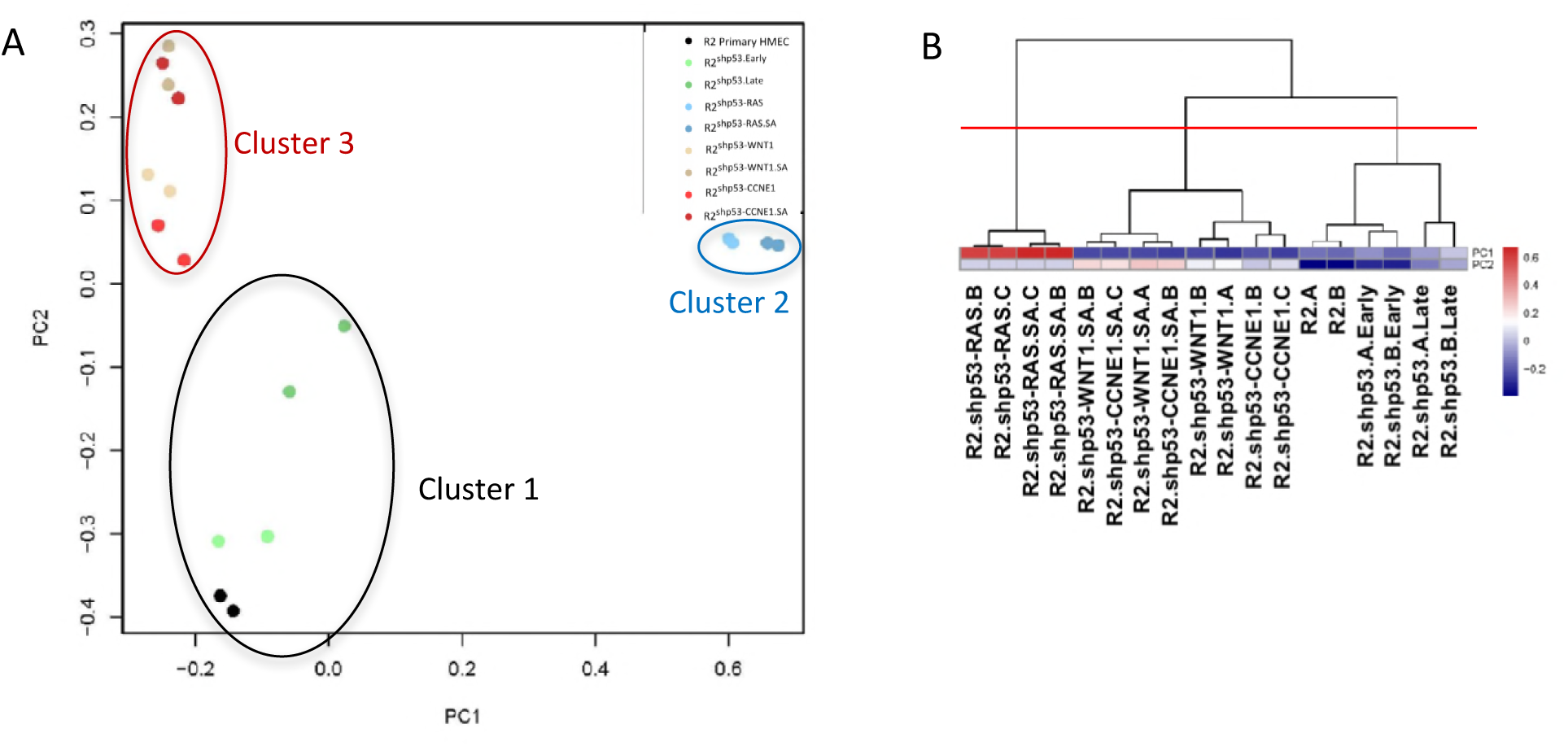
A. Joint PCA analysis including CNA, DNA methylation, miR and mRNA data B. Non supervised hierarchical clustering (Ward) on PC1 and PC2 segregating in three clusters (red-line threshold)

### Immortalized and transformed HMECs form 3 distinct clusters defined by the activated oncogene

Next, we determined whether the different immortalized and transformed HMEC sublines presented distinct profiles of genetic and epigenetic anomalies. Copy Number Alterations (CNA), DNA Methylation, miRNA (miR) and mRNA expression levels were determined at different steps of cell immortalization and transformation. To integrate these data corresponding to diverse molecular features and different technological platforms and increase the possibility of discovering coordinated changes, we used moCluster (moGSA package), a multivariate single sample gene-set analysis method, developed to produce integrative clustering on multiple omics data. In brief, this approach is based on multivariate latent variable decomposition to discover correlated global variance structure across datasets [19].

Two independent biological replicates of each immortalization or transformation step, (R2, R2^shp53^, R2^shp53-WNT1^, R2^shp53-CCNE1^, R2^shp53-RAS^ and R2^shp53-WNT1.SA^, R2^shp53-CCNE1.SA^, R2^shp53-RAS.SA^ HMECs) were included in this analysis. These analyses pointed at similarities and differences between distinct sublines and stages of transformation. These were determined by means of Principal Component Analysis (PCA). Changes defining the most variant PCA vector were subsequently analyzed by Ward clustering. The combined analysis of the four datasets (CNA, DNA methylation, mRNA and miRNA expression) defined 3 clusters. The first one, at the trunk, organized around R2 and R2^shp53^ early and late passages. The second corresponded to the four R2^shp53-RAS^ sublines. The third cluster included R2^shp53-WNT1^ and R2^shp53-CCNE1^ sublines (Fig 2A). The number of clusters was confirmed by Nbclust (Fig 2B and S Appendix for details). Hence, the MoGSA analysis revealed that the HMEC sublines transduced with different genetic elements showed clear differences in their genetic and epigenetic patterns. We next analyzed CNAs, DNA methylation, mRNA and miRNA datasets individually in order to verify whether the differences in patterns applied similarly to all datasets.

### Overexpression of CCNE1, WNT1 or HRAS^v12^ oncogenes in immortalized HMECs result in distinct profiles of copy number alterations

Genomic regions involved in CNAs were identified with the Nexus 7.5 Software (Biodiscovery, CA., USA) and only CNAs covering at least 2 Mb were used in the moGSA analysis. The PCA classification of the different HMEC sublines singled out three clusters. Cluster 1 positioned at the trunk encompassed R2 and R2*^shp53^* replicates, cluster 2 formed by the R2^shp53-RAS^ and R2^shp53-RAS.SA^ sublines and a more dispersed cluster 3 comprising R2^shp53-WNT1^, R2^shp53-CCNE1^, R2^shp53-WNT1.SA^ and R2^shp53-CCNE1.SA^ (Fig 3A). The distance along the PC1 vector in the principal component analysis, separating cluster 2 from cluster 3, illustrated the strong differences at the CNA level between the RAS and the WNT1 or CCNE1 transformed sublines (Fig 3A). A clustering analysis of the CNAs defining the PC1 vector identified co-occurring loss of chromosome 4, loss at 8p and gain at 8q as the most significant anomalies characterizing the R2^shp53-RAS^ HMECs (Fig 3B). Other events were gains at 12q and losses at 14 and 18q (S4A-B Fig). In clear contrast, R2^shp53-WNT1^ and R2^shp53-CCNE1^ HMECs were characterized by losses at chromosomes 2, 3 and 6, as well as focal gains at 11q13 and 20q13 (Fig 3B and S4A-B Fig). Interestingly, whereas the pre-transformed R2*^shp53-RAS^* showed little difference with their transformed R2^shp53-RAS.SA^ counterpart, significant deviation was detected between R2^shp53-CCNE1^ and R2^shp53-CCNE1.SA^ as well as between R2^shp53-WNT1^ and R2^shp53-WNT1.SA^, with additional gains at chromosome 11q13-q14 and 20q11-q13, respectively (Fig 3B). Together with the global increase in the fraction of the genome involved in CNAs, these data suggested an increase of genetic instability in R2^shp53-CCNE1.SA^ and R2^shp53-WNT1.SA^ (Fig 1D). In contrast, R2^shp53-RAS^ HMECs did not show significant changes in CNAs after soft agar cloning (Fig 1D and Fig 3B).

**Fig. 3:**
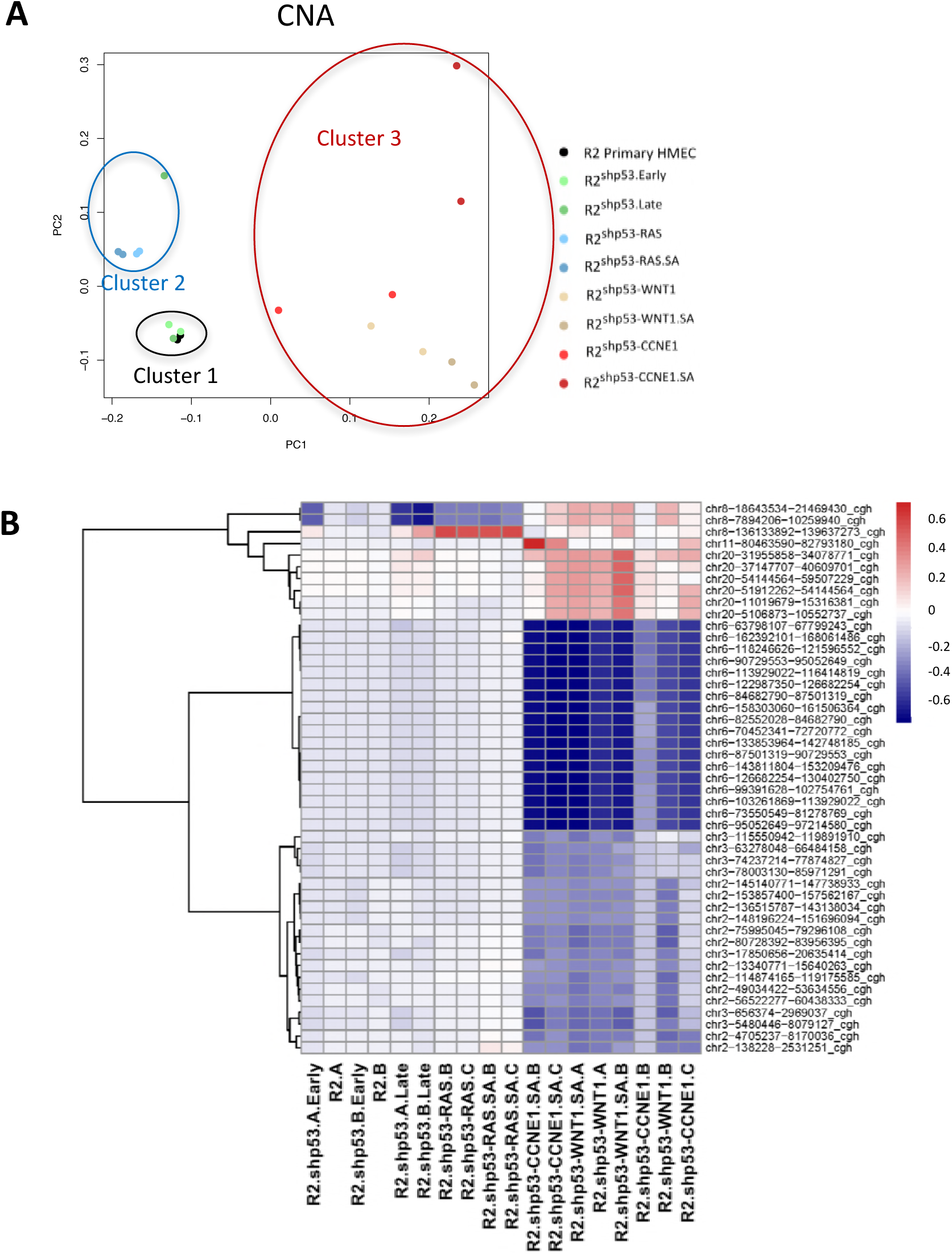
Copy Number Alterations profiles vary according to the oncogene expressed. Copy Number Change analyses were performed on HG18 CGH 385K Whole Genome v2.0 array. Regions of gain or loss were determined using the Nexus 7.5 Software and intervals of at least 2 Mb selected. **A.:** PCA analysis of the CNAs. **B:** Ward clustering of the CNAs defining PC1 in the PCA analysis, color code of the heatmap is blue for loss, red for gain.

### Overexpression of CCNE1, WNT1 or HRAS^v12^ oncogenes in immortalized HMECs result in distinct profiles of DNA methylation

DNA methylation profiles of the HMEC sublines were analyzed at the single-base resolution level by reduced representation bisulfite sequencing (RRBS). Variations in methylation levels were quantified at sequences containing at least 5 contiguous CpG pairs genome-wide. A global increase and change in distribution of the methylation pattern was clearly detected between primary R2 HMECs and all the sublines derived there from (Fig 4A). Variations were particularly pronounced in transformed cells after soft agar cloning (R2^shp53-RAS.SA^, R2^shp53-CCNE1.SA^ and R2^shp53-WNT1.SA^). This increase in CpG methylation suggested a global DNA methylation reprogramming in shp53 immortalized and oncogene transformed HMECs. Using a methylation difference of 0.2 as inclusion criterion, we next performed a SAMR analysis of these data to identify differentially methylated regions (DMRs) that varied most significantly between sublines. Identified DMRs were analyzed in moGSA and used to classify cell lines by Principal Component Analysis (PCA). The PCA classification based on DNA methylation was remarkably similar to that generated with CNAs. Here again R2^shp53-RAS^ and R2^shp53-RAS.SA^ defined a clearly distinct cluster of the R2^shp53-CCNE1^, R2^shp53-WNT1^, R2^shp53-CCNE1.SA^ and R2^shp53-WNT1.SA^ luster (Fig 4B). Additional clustering analysis of the DMRs centered on CpGs methylation patterns, confirmed this trend, R2^shp53-RAS^ HMECs showing an almost binary difference when compared to R2^shp53-CCNE1^ and R2^shp53-WNT1^ (Fig 4C). Noticeably, primary R2 and immortalized R2^shp53^ (Early and Late passages) clustered at the trunk (cluster 1 in Fig 4B).

**Fig. 4:**
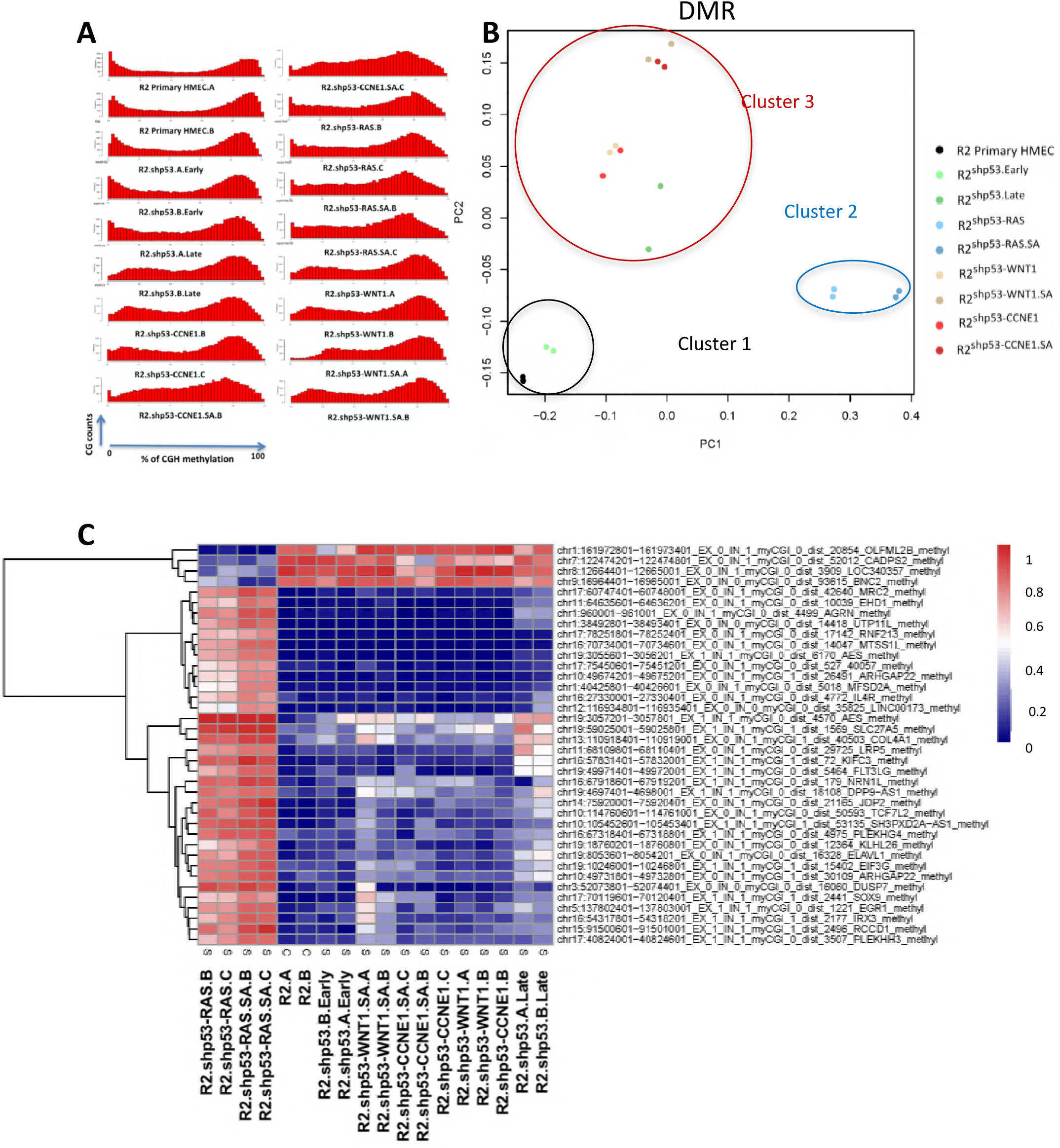
CpG methylation profiles show coordinated variation according to the oncogene expressed. **A.** density histograms of RRBS methylation scores at CpGs sites (at least 5 contiguous CG) genome wide in primary R2 HMECs, shp53.Early, shp53.Late, shp53-CCNE1, shp53-CCNE1.SA (Soft Agar), shp53-WNT1, shp53-WNT1.SA, shp53-RAS, shp53-RAS.SA. **B.** PCA analysis of the most significantly varying DMRs. **C;** Ward clustering of the DMRs (without constraint concerning their location) defining PC1 in the PCA analysis, color code of the heatmap is red for hypermethylation, blue for hypomethylation.

Altogether, both CNA and DNA methylation profiles indicated that the ectopic expression of an oncogenic form of RAS in R2*^shp53^* HMECs induced specific genomic and epigenetic alterations, that markedly differed from those observed in HMECs transduced with *WNT1* or *CCNE1*. Interestingly, in the same analyses WNT1 or CCNE1 transformed HMECs co-clustered in both classifications, suggesting these two oncogenic events result in globally similar adaptive programs.

### Overexpression of CCNE1, WNT1, or HaRAS^v12^ oncogenes in R2^shp53^ HMECs result in distinct transcriptional programs and phenotypical cell fates

We next analyzed the mRNA and miRNA expression profiles. As observed with CNA and DNA methylation profiles, cluster analyses of both miR and mRNA expression profiles clearly distinguished the RAS and CCNE1/WNT1 transformed cells in two separate clusters (Fig 5A and Fig 5C). We observed that many genes presented an inverted pattern of expression in the RAS compared to the CCNE1/WNT1 sublines (Fig 5B and Fig 5D) in the Ward clustering analysis of expression changes defined by the PC1 vector in the miR and mRNA datasets. GeneGo Metacore (Thompson Reuter) network analyses of the miR and mRNA gene lists that discriminated cluster 2 and cluster 3 identified several differentially activated functional gene networks. Based on the miR dataset, this analysis clearly pointed at strong differences in EMT-associated pathways (S5 Fig), illustrated by the level of miR200c, a well characterized regulator of the expression of the EMT transcription factor ZEB1 [20], and the mesenchymal cell-specific miR143/miR145, that differed markedly between RAS and CCNE1/WNT1 cells [21]. Morphological features (cuboidal *vs.* fusiform) and immunofluorescence staining of cells for the epithelial E-Cadherin (ECAD) and the mesenchymal Vimentin (VIM) markers, confirmed that primary R2, immortalized R2*^shp53^*, and transformed R2*^shp53-CCNE1^* and R2*^shp53-WNT1^* cells all maintained an epithelial phenotype (cuboid, ECAD^high^, VIM^low^), while R2*^shp53-RAS^* cells were fusiform, ECAD-negative and VIM^high^ indicating a clear mesenchymal conversion (S6 Fig).

**Fig. 5:**
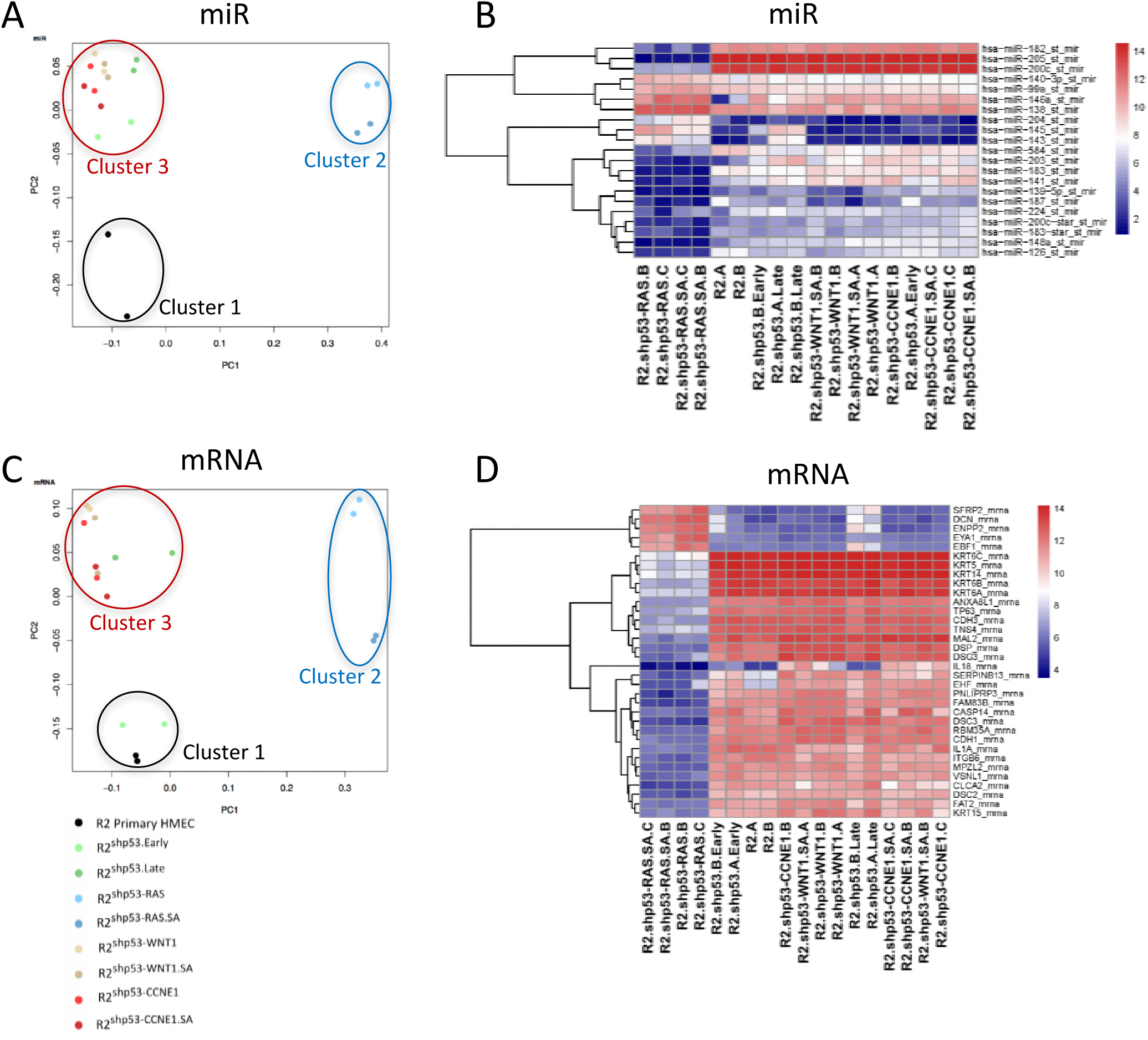
miR and mRNA expression profiles of the shp53-RAS show clear differences from those of shp53-CCNE1 and shp53-WNT1 sublines. A: PCA analysis of the most significantly varying miR. B: Ward clustering of the miR defining PC1 in the PCA analysis, color code of the heatmap is red for overexpression, blue for underexpression. C: PCA analysis of the most significantly varying mRNA. (D) Ward clustering of the miRdefining PC1 in the PCA analysis, color code of the heatmap is red for overexpression, blue for underexpression.

### Chronology of the genetic and epigenetic changes occurring during transformation of HMECs

Monitoring the genetic and epigenetic changes occurring at each step of the cell transformation process allowed us to reconstruct their chronology. This could be inferred from the PC2 vectors in the PCA classifications. Starting from normal R2 primary cells, moving to early and late immortalized cells and then to anchorage independent cells growing in soft agar, we noted that the different sublines bore different positions in these classifications according to the dataset considered. In the miR classification, the immortalized R2^shp53^ (Early and Late passages) were separated from primary R2 and co-clustered with R2^shp53-CCNE1^ and R2^shp53-WNT1^ (Fig 5A). In contrast, in the mRNA-based and DMR classifications, primary R2 co-clustered with early passage R2^shp53^ cells. Late passage R2*^shp53^* had undergone changes that brought them closer to R2^shp53-CCNE1^ and R2^shp53-WNT1^ (Fig 4B and Fig 5C). Finally, the classification based on CNAs, revealed that early and late passages R2^shp53^ co-clustered with R2 (Fig 2B). These co-clustering analyses suggested a temporal hierarchy of the genetic and epigenetic events that primed transformation in our models. The clear clustering differences observed between early and late passage R2^shp53^ cells according to either the miR, mRNA, DMR or CNA datasets, strongly suggested that upon p53 inactivation in HMECs, miR expression was the first to be modified, whilst the modification of mRNA expression profiles and massive modifications of the methylation landscape occurred later. Upon transduction with RAS, WNT1 or CCNE1, additional DNA methylation changes occurred concomitantly with modifications in mRNA expression. Copy number changes emerged last and underwent a clear selection process during soft agar cloning (Fig 2B and S4A Fig).

### Impact of copy number changes and differential DNA methylation on gene expression

First, we identified 2182 genes whose expression change was correlated to CNAs and in a second time selected the genes that were differentially expressed in the RAS or the CCNE1/WNT1 clusters compared to R2 and R2^shp53^ HMECs. We noted that genes with copy number dependent expression change in the R2^shp53-RAS^ showed very little overlap with those modified in R2^shp53-CCNE1^/R2^shp53-WNT1^, since only 2 (1.7%) genes were present in both lists (S7A Fig). This absence of significant overlap could not have happened by chance (hypergeometric test, p=0.75). In the R2^shp53-RAS,^ we identified 90 genes (36 overexpressed in regions of gain and 54 underexpressed in regions of loss) differentially expressed relative to R2 and R2^shp53^ HMECs. Similarly, 156 genes (118 overexpressed in regions of gain, 38 underexpressed in regions of loss) were differentially expressed in the R2^shp53-CCNE1^/R2^shp53-WNT1^ cluster compared to R2 and R2^shp53^. A large proportion of these genes mapped in the CNA regions that were most discriminating to either R2^shp53-RAS^ (65% in the gain at chromosome 8 or loss at chromosome 4) or to R2^shp53-CCNE1^/R2^shp53-WNT1^ (58% in the gains at 8p21, 11q13, 17q21 or 20q11-q13, losses at 6q) (S1 Table). We also noted that 78 genes, gained and overexpressed in either R2^shp53-RAS^ or R2^shp53-CCNE1^/R2^shp53-WNT1^ HMECs were part of the gene list previously reported to be associated with common regions of amplification in breast cancer [22]. This enrichment was highly significant (hypergeometric test p= 1.10^-13^) and suggested that our HMEC stepwise transformation models shared genetic pathways with breast cancer.

We selected the genes annotated as cancer genes (oncogenes and tumor suppressors) and for R2^shp53-RAS^ HMECs added genes belonging to the RAS pathway (Fig 6A and S1 Table). In the R2^shp53-RAS^ cluster, this revealed 5 gained and over-expressed (*ERBB2, HEY1, PLAG1, ZBTB10, CRTC3*), all known oncogenes, and 6 lost and under-expressed genes. Of the 6 lost and under-expressed genes, 4 were components of a RAS pathway (*TMEM154, LRAT, AKAP6, MTUS1*) and 2 were cancer suppressor genes (*FBXW7, SERPINB5*). *FBXW7* encodes an ubiquitin ligase that regulates negatively transcription factors such as MYC, FOS or NOTCH [23], whereas *SERPINB5 (MASPIN)* is described as a metastasis suppressor in breast and other cancers [24].

**Fig. 6:**
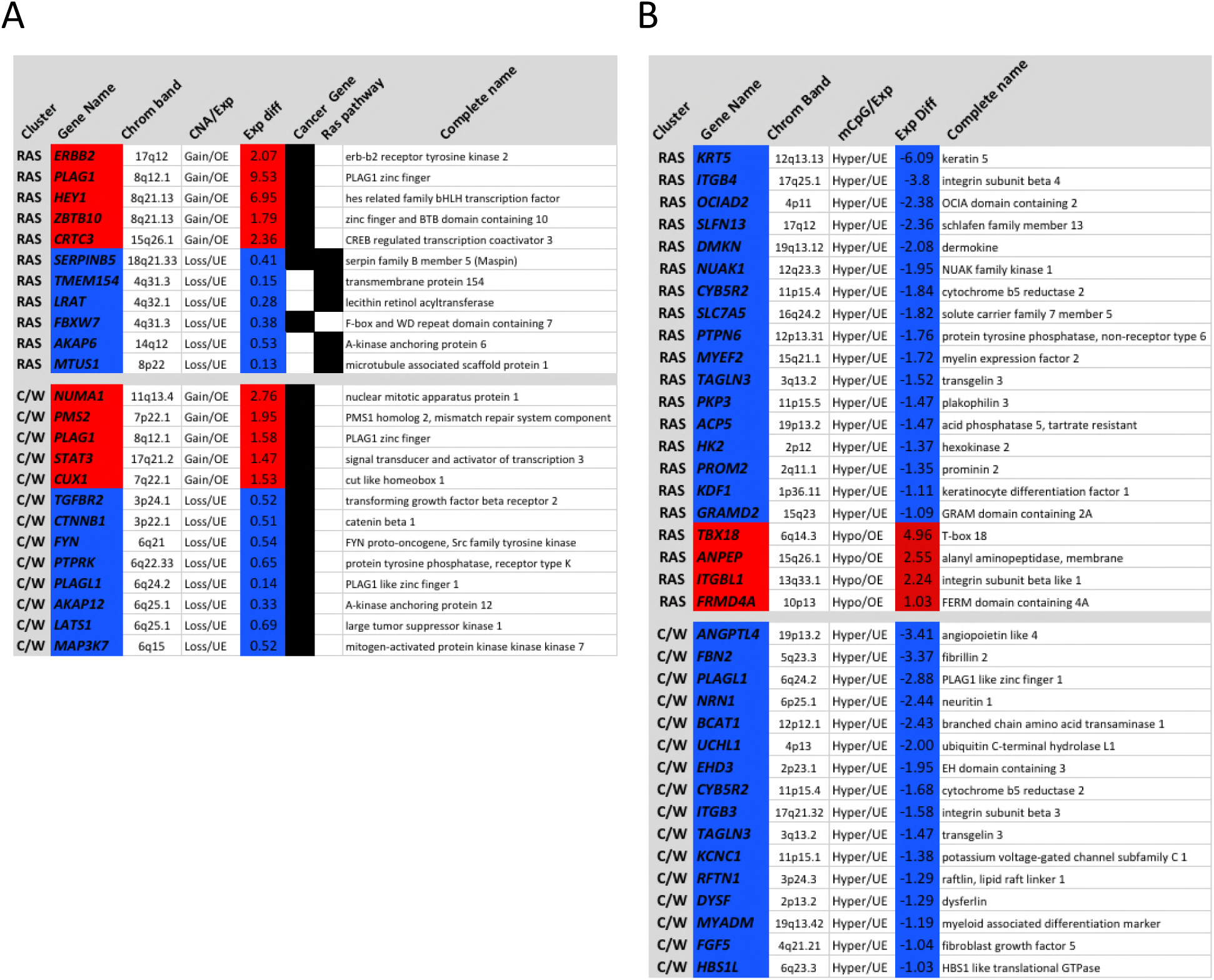
Most significant gene expression differences due to copy number alterations (A) and differential methylation (B) in R2^shp53-RAS^ and R2^shp53-CCNE1^/R2^shp53-WNT1^ HMECs. Expression difference levels (Exp diff column) have been calculated relative to expression levels in the R2 and R2^shp53^ cluster and indicated in log2 scale. Genes are listed for each cluster (RAS for R2^shp53-RAS^, C/W for R2^shp53-CCNE1^/R2^shp53-WNT1^). A: Genes modified as a consequence of CNA have been selected from a broader list on the basis of their assignment either as cancer or tumor suppressor genes (assembled under the cancer gene category) or as members of the RAS pathway. Highlighted in red, genes overexpressed in regions of gain (Gain/OE), in blue, genes underexpressed in regions of loss (Loss/UE). B: Genes modified as a consequence of differential methylation at the transcription start sites (TSS). In red, genes hypomethylated with increased expression (Hypo/OE), in blue, genes hypermethylated with reduced expression (Hyper/UE).

In the R2^shp53-CCNE1^/R2^shp53-WNT1^ cluster, we identified 13 genes, 5 gained and over-expressed, 8 lost and under-expressed (Fig 6A). All 13 genes classified as cancer genes. Of the 5 gained and over-expressed genes, *NUMA1*, *PMS2* and *CUX1* have well documented roles in mitotic spindle assembly or DNA repair and *STAT3* is a key signaling node for a number of growth factors and cytokines frequently involved in cancer [25]. *PLAG1* was selected both in the RAS and CCNE1/WNT1 upregulated genes, but its expression level was 9 time higher in R2^shp53-RAS^ compared to R2^shp53-CCNE1^/R2^shp53-WNT1^. Of the 8 lost and under-expressed genes in the CCNE1/WNT1 cluster, we noted *LATS1,* negative regulator of the Hippo pathway, *CTNNB1* (β*-*catenin), the *SRC* homolog *FYN* and *TGFBR2*.

Using a similar approach, we also searched for genes whose expression was impacted by CpG methylation. We restricted our analysis to DMRs located close to transcription start sites (TSS) of annotated genes showing at least a 20% change in methylation, associated to a 2fold variation in mRNA expression level (Fig 6B). Again, genes modified by DNA methylation in the R2^shp53-RAS^ cluster showed little overlap with those in R2^shp53-CCNE1^/R2^shp53-WNT1^ HMECs (Hypergeometric test, p =0.66, S7B Fig). In the R2^shp53-RAS^ cluster, 17 genes were hypermethylated and underexpressed, 7 of which were key in cell-adhesion and the epithelial phenotype (*KRT5, ITGB4, DMKN, PKP3, ACP, PROM2, KDF1*). Interestingly, we identified 4 hypomethylated and overexpressed genes, of which 2 (*ITGBL1, FRMD4A*) corresponded to genes involved in cell invasion and metastasis [26,27]. In the R2^shp53-CCNE1^/R2^shp53-WNT1^ HMECs, 16 genes all hypermethylated and downregulated were identified (Fig 6B). The most salient feature was the downregulation of genes related to cell invasion or EMT (*ANGPTL4, ITGB3, EHD3, FBN2*). Interestingly, the products of two of these genes, *ITGB3* and *EHD3,* interact physically in an activating regulatory loop.

Altogether, the RAS and the CCNE1/WNT1 transformed HMECs showed clear differences in genes whose expression was impacted by either CNAs or CpG methylation. The principal features in RAS HMECs were the repression of genes associated with the epithelial phenotype and cell adhesion and conversely the activation of genes favoring cell invasion. In clear contrast, CCNE1/WNT1 HMECs showed a downregulation of EMT or invasion associated genes, combined with the activation of DNA repair and cell division genes. These results indicate the activation of distinct pathways in the respective sublines, with an opposite trend concerning cell phenotype, pro-mesenchymal in R2^shp53-RAS^ and pro-epithelial in R2^shp53-CCNE1^/R2^shp53-WNT1^.

### RAS HMECs and CCNE1/WNT1 HMECs resemble claudin-low and Basal-like breast cancer, respectively

We wanted to determine how much our HMEC models paralleled with human breast cancer. We first noted that the 16 HMEC models were classified as Basal-like breast cancer according to the CIT [9] or the BasalA/B [28] breast cancer classifiers (S8A Fig). Using a restricted Claudin-low classifier [29], R2*^shp53-RAS^* were classified as Claudin-low, R2*^shp53-CCNE1^* and R2*^shp53-WNT1^* as epithelial, while R2 and R2*^shp53^* formed an intermediate group, combining characteristics of both the Claudin-low and the epithelial group (Fig 7A). Next, we sought to compare the characteristics of our HMEC models with that of the Claudin-low and the Basal-like breast cancer subgroups in the human primary breast cancer METABRIC dataset. Restricting our analysis to *TP53* mutated and Basal-like tumors (defined with the PAM50 classifier), we selected 259 tumors, which we stratified in Claudin-low/Basal-like (83) and Basal-like/Non-Claudin-low (176) subsets. We determined that tumors in this subgroup presented a significantly lower incidence of CNAs than the Basal-like/Non-Claudin-low (Fig 7B-C) and that the KRAS pathway was among the top activated pathways in the Claudin-low/Basal-like tumors (S9A-B Fig). In addition, *CCNE1* overexpression was significantly more frequent in the Basal-like/Non-Claudin-low than in the Claudin-low subgroup (Fig 7D). These data suggested that the differences in genomic profiles induced by distinct oncogenes observed in our HMEC models mimicked situations occurring in spontaneous human breast tumors: fewer genomic changes and elevated RAS in mesenchymal like claudin-low basal tumors, in comparison to non-claudin-low basal tumors which show frequent *CCNE1* overexpression and an epithelial phenotype.

**Fig. 7:**
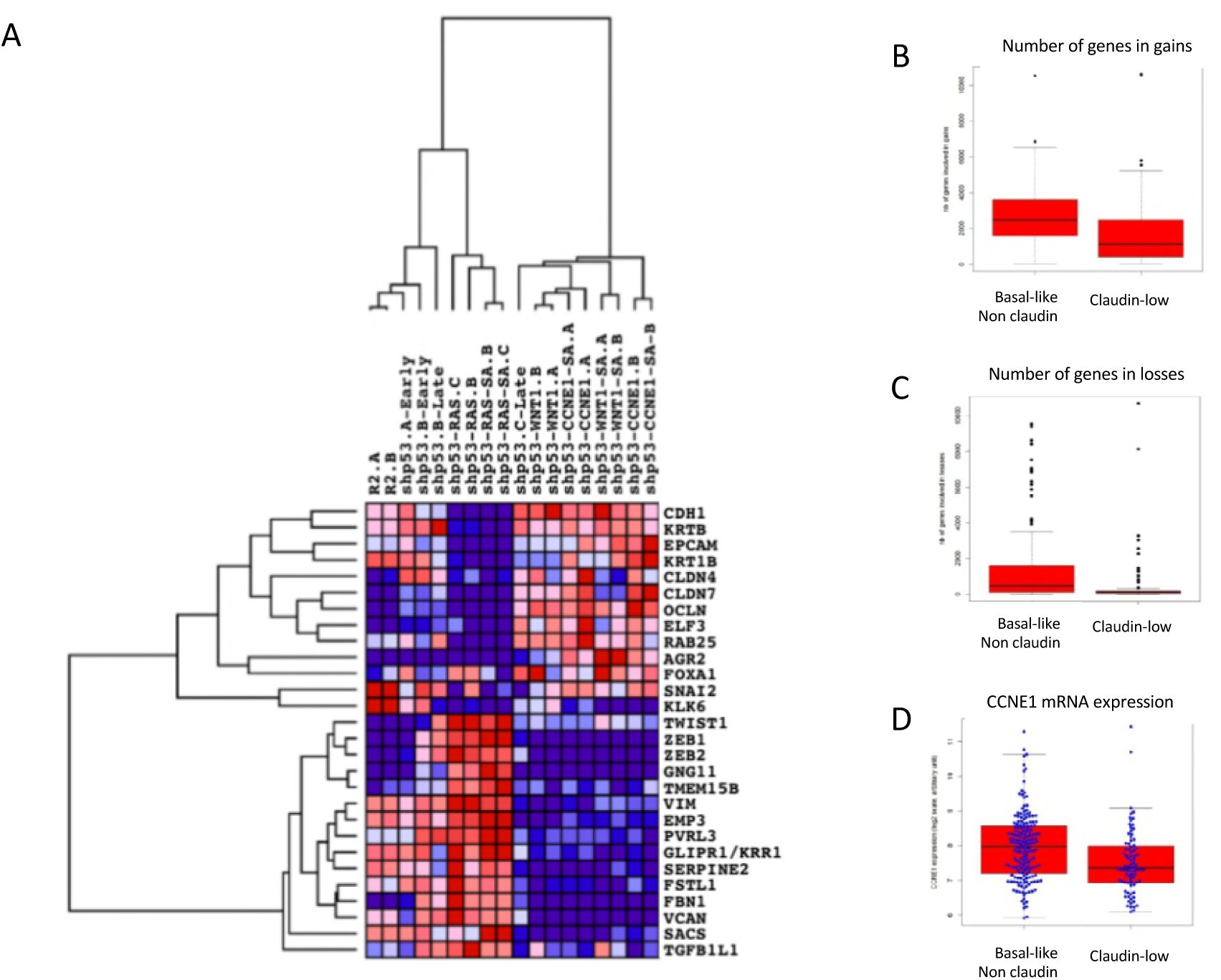
The different HMEC models resemble basal breast cancers and show varying levels of mensenchymal traits. **A.** clustering analysis using a 28 gene signature representative of claudin low tumors. R2shp53-RAS models classified as strict claudin-low (mesenchymal), whereas R2shp53-CCNE1 and R2shp53-WNT1 were strictly epithelial. Of note, R2 (Ctrl here) and R2shp53 presented intermediate profiles combining elevated expression of both epithelial and mesenchymal genes. **B-C.**CNE1/WNT1 and RAS HMECs resemble respectively Basal-like and claudin-low breast cancers whichshow similar differences in CNA numbers, **D.** differences in CCNE1 mRNA expression levels. This analysis was done on 259 Basal-like tumors (PAM 50 classification) selected from the Metabric dataset which were split in Basal-Like/Non claudin (176) and Claudin-low (83).

## Discussion

Breast cancer can be broken down into 5 to 12 molecular subtypes on the basis of differences in mRNA expression and genetic anomaly profiles [9,10]. Although direct evidence is scarce, it is generally assumed that the origin of these molecular subtypes lies in cell lineage differences, where the original tumorigenic insult took place [8,9]. To explain why premenopausal breast cancers are frequently of the triple negative type, whereas ER+ luminal type tumors are prevalent in post-menopausal patients, it has been proposed that the mammary gland undergoes an aging related cell lineage drift from the basal/myoepithelial to the luminal lineage which becomes predominant after menopause [30,31]. On the other hand, luminal progenitor cells have been shown to be at the origin of basal-like breast tumors in patients with constitutional *BRCA1* inactivation [32,33]. The fact that *BRCA1* mutations induced a differentiation shift from luminal progenitors towards a basal-like rather than a luminal phenotype helped explain this apparent contradiction [34]. Similarly, expression of an activated *PikcaH1047R* allele in committed unipotent luminal cells in mouse mammary glands induced cell fate reprogramming and emergence of basal like tumors [35]. These data are thus in favor of the idea that founding oncogenic mutations could impinge on tumor subtype in breast cancer. This is further supported by the work of Ben-David and coauthors (2016) [36] showing that, in genetically modified mouse mammary tumors, CNA profiles differed according to the driver mutation initiating the tumor. Using primary HMECs, which according to the classification proposed by Lim and coworkers (2009) [32] correspond to luminal progenitors (S8B Fig), we show here that cell transformation by way of distinct oncogenes resulted in different patterns of aberrations at both the CNA and DNA methylation levels. Most remarkably, HMECs transformed by RASv12 (R2^shp53-RAS^) presented clearly distinct patterns of genetic and epigenetic modifications compared to their R2^shp53-CCNE1^ or R2^shp53-WNT1^ counterparts. The latter two exhibited globally similar CNA and DNA methylation profiles, albeit some focal differences could be found.

In our model system, inactivation of the *TP53* gene was the initial step towards transformation. Given the role of *TP53* in keeping the integrity of the genome, onset of genetic instability in R2^shp53^ HMECs was expectable. Surprisingly, no gross genomic anomalies, even after more than one year in culture, was observed. R2^shp53^ rapidly became tetraploid upon inactivation of p53, in line with the critical role of p53 in ploidy control [37]. Tetraploidy is considered as a prelude to large scale chromosomal aberrations in cancer [38-40] and has been shown to confer rapid adaptation capacity in yeast [41]. We thus propose that while p53 inactivation did not provoke marked genetic changes in our HMEC models, it favored genetic plasticity and laid the ground for further genetic rearrangements upon oncogene expression. This trend was illustrated by R2^shp53-CCNE1^ and R2^shp53-WNT1^, which progressively became aneuploid and whose rearrangement levels increased significantly after soft agar cloning. In remarkable contrast, R2^shp53-RAS^ remained strictly tetraploid and did not acquire further anomalies after soft agar. This suggested ongoing genetic instability in R2^shp53-CCNE1^ and R2^shp53-WNT1^ HMECs, whereas a stabilization took place in R2^shp53-RAS^. This difference in genetic instability levels between the RAS and the CCNE1 or WNT1 transformed HMECs was further supported by the reduced number of γH2Ax and 53BP1 foci in R2^shp53-RAS^ compared to R2^shp53-CCNE1^ or R2^shp53-WNT1^. These observations are in line with recent work showing that primary mammary epithelial cells or immortalized HME expressing high levels of *ZEB1* kept stable genomes upon transduction of RASv12 [42]. In this model system, *ZEB1* was shown to control a ROS scavenging program that protected cells overexpressing RASv12 from DNA damage. The relative genetic stability of R2^shp53-RAS^ HMECs could be linked to the strong EMT, associated with the activation of *ZEB1,* that characterized these cells (Fig 7A and S8B Fig). In contrast, R2^shp53-CCNE1^ and R2^shp53-WNT1^ HMECs showed an epithelial phenotype and low levels of *ZEB1* expression. R2^shp53-RAS^ were Claudin-low, whereas R2^shp53-CCNE1^ classified and R2^shp53-WNT1^ classified as Basal-like with epithelial dominance. Interestingly, R2^shp53^ cells formed an intermediary cluster combining epithelial and mesenchymal features (Fig 7A). This suggested that the phenotypic switch (EMT for R2^shp53-RAS^ and MET for R2^shp53-CCNE1^ and R2^shp53-WNT1)^ occurred as a consequence of the ectopic expression of the respective oncogenes. Remarkably, the R2^shp53-RAS^ resembled Claudin-low breast cancer, whereas R2^shp53-CCNE1^ and R2^shp53-WNT1^ were analogous to basal-like breast tumors. As a matter of fact, Claudin-low breast tumors showed a lower level of CNAs and a frequent activation of the RAS pathway, whereas basal-like tumors presented higher CNA levels and elevated *CCNE1* expression. Although our HMEC models do not sum up the complete spectrum of human breast cancers, our data show that they mimic at least partially Basal-like and Claudin-low breast cancer.

In this work, we inferred a chronology of the genetic changes that occurred at different steps of HMEC transformation. The first modified were miR and mRNA expression, followed closely by DNA methylation and CNAs occurred last. This indicated that expression changes acted as drivers, modified the phenotype and impacted on the epigenetic and genetic landscape that finally locked the changes. In R2^shp53-RAS^ HMECs, a large number of genes that were modified by DNA methylation were involved in EMT, such as the repression of the epithelial genes *KRT5*, *ITGB4*, *NUAK1*, *ACP5* and *PROM2,* concomitant to the hypomethylation and overexpression of the pro-invasive *ITGBL1* and *FRMD4A* genes. DNA methylation changes have been proposed as hallmarks of advanced stages of EMT [43]. Thus, epigenetic changes in R2^shp53-RAS^ are in coherence with the deep shift towards a mesenchymal phenotype undergone by these cells. Interestingly, CNAs in R2^shp53-RAS^ impacted two actors of the NOTCH pathway, with *HEY1* being gained and overexpressed and *FBXW7* being lost and downregulated. *HEY1* is an important NOTCH target gene and *FBXW7* is a repressor of NOTCH [23]. Interestingly, FBXW7 ubiquitination activity has been shown to be repressed by activated RAS [44]. Thus, the copy number reduction of *FBXW7* could be an adaptive response to RAS activation, resulting in the activation of the NOTCH pathway in R2^shp53-RAS.^ In R2^shp53-CCNE1^/R2^shp53-WNT1^. We noted hypermethylation and downregulation of the *ANGPTL4, ITGB3, EHD3* and *FBN2* genes, which have been associated to cell migration and invasion. Hence, in R2^shp53-CCNE1^/R2^shp53-WNT1^ DNA methylation changes contributed to the mesenchymal to epithelial transition demonstrated by the mRNA expression changes. This trend was reinforced by CNAs, as shown by the loss and downregulation of *TGFBR2* and *CTNNB1*, indicating the downregulation of the TGFβ pathway in CCNE1 or WNT1 HMECs. The TGFβ pathway is a known promoter of EMT and its downregulation appeared in coherence with the epithelial phenotype of R2^shp53-CCNE1^ and R2^shp53-WNT1^ cells. Further notable consequence of CNAs in R2^shp53-CCNE1^/R2^shp53-WNT1^ cells were the gain of the mitotic spindle assembly genes *NUMA1, CUX1* or mitosis and loss of *AKAP12* negative regulator of Polo Kinase,. These anomalies could be associated to aneuploidy in these cells. Indeed, depletion of *AKAP12* has been shown to lead to aneuploidy and increased tumor growth [45] and overexpression of *NUMA1* and *CUX1* could represent adaptive responses to increasing chromosomal instability.

## Conclusions

Altogether, data presented herein show that R2^shp53-RAS^ HMECs presented clearly different genomic and DNA methylation profiles relative to R2^shp53-CCNE1^/R2^shp53-WNT1^. The nature of the genes whose expression was modified either by DNA methylation or CNAs were consistent with the dominant pathways activated and reflected the phenotypes in the respective models, mesenchymal in R2^shp53-RAS,^ epithelial in R2^shp53-CCNE1^/R2^shp53-WNT1^ (Fig 8).

**Fig 8:**
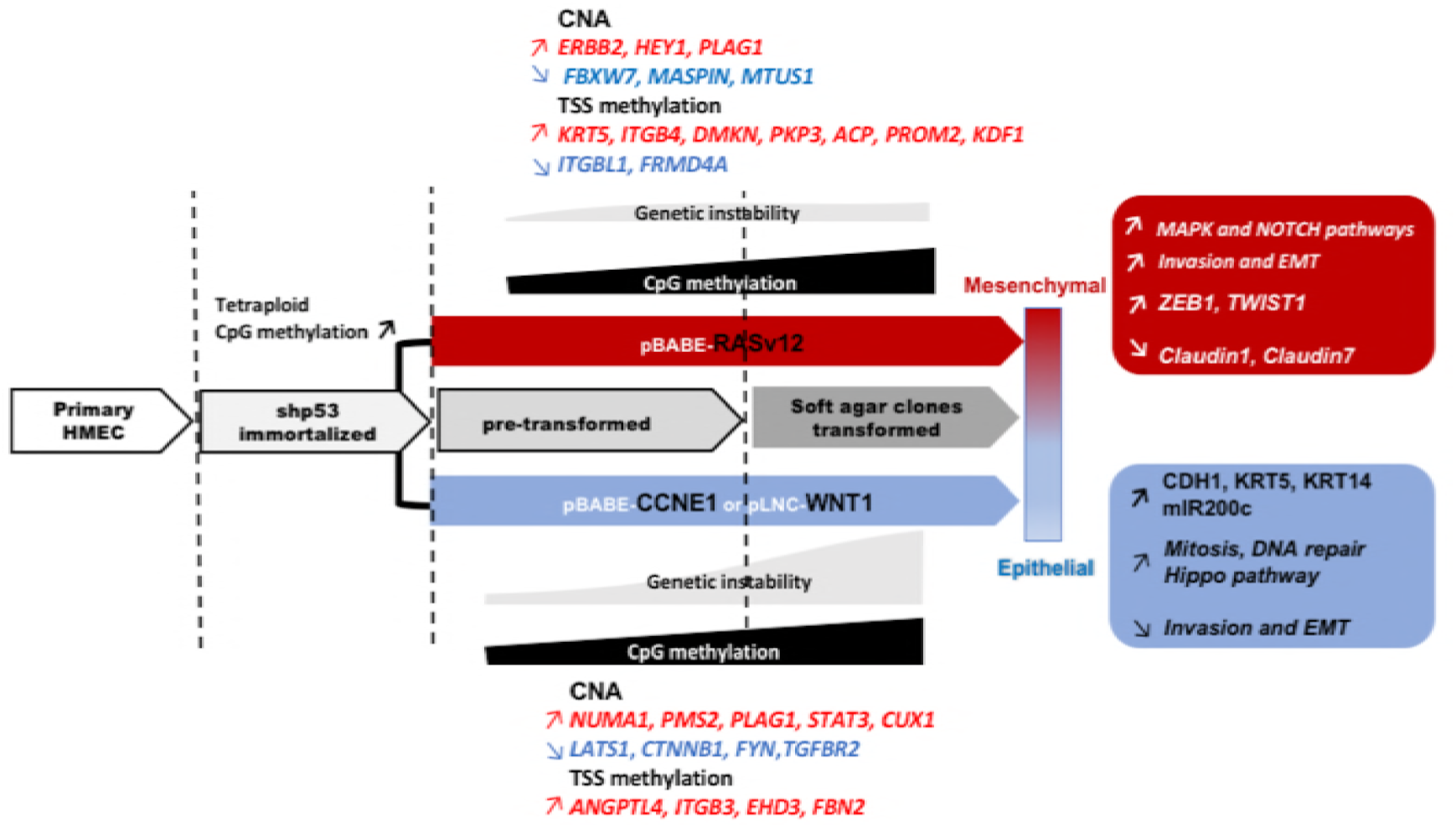
Genetic, epigenetic and phenotypic modifications resulting from the transduction of distinct oncogenes in HMEC. The changes occurring are depicted at different steps starting from primary HMEC and ending at transformed cells isolated from soft agar colonies. For readability purposes, only principal events and genes are presented. Arrows indicate upregulation or downregulation of expression of genes modified by CNAs or differential methylation at their TSS, as well as up or downregulation of pathways as part of the phenotypic consequences of the activation of the respective oncogenes.

Thus, our data strongly suggest that early activation of distinct oncogenic insults in a given cell type will not only impinge on the phenotypic characteristics of the resulting tumors, but also impact on their genomic and epigenetic landscapes and could contribute to the determination of cancer subtypes.

## Material and Methods

### HMEC models

Human Mammary Epithelial Cells (HMECs) were isolated from mammary gland explants obtained from plastic surgical after informed consent from the patient. This work was approved by the Ethics committee of the University of Montpellier. Cell suspensions were produced by mechanical and enzymatic dissociation with 1% collagenase. After elimination of fibroblasts, HMECs were cultured at 37°C in 5% CO2, in conditioned MEBM medium supplemented with antibiotics (*MEGM single Quots*, Lonza, Levallois-Perret, France). Primary HMECs were transduced with amphotropic retroviral supernatants corresponding to: pSUPER.retro.hygro-shp53, pBABE.neo-CCNE1, pLNC-WNT1, pBABE.puro-HRASV12 followed by 3 weeks of antibiotic selection.

### β-galactosidase senescence test

**β**-galactosidase activity was assay by histochemistry using the *Senescence Cells Histochemical Staining* kit (Sigma-Aldrich, St Quentin Fallavier, France) following manufacturer’s instructions. **β**-galactosidase positive cells were quantified under the microscope in duplicate on 400 cells minimum.

### Telomeric restriction fragment (TRF) analysis and telomerase activity test and chromosome counts

TRF were purified using the *TeloTAGGG Telomere Length Assay* (Sigma-Aldrich, St Quentin Fallavier, France) following manufacturer’s instructions. TRF sizes determined by 0.8% agarose gel electrophoresis and Southern blotting. The Nylon membrane was hybridized telomeric DNA probe labeled with digoxigenine and revealed with anti-DIG antibodies coupled alcalin phosphatase and chemioluminescence. Medium TRF size was calculated using TRF=Σ(ODi)/Σ(ODi/Li) (ODi= optical density at i, Li= size at i) Telomerase activity was measured on 2×10^5^ cells with the *TeloTAGGG Telomerase PCR ELISA* kit (Sigma-Aldrich, St Quentin Fallavier, France) following manufacturer’s instructions.

### Anchorage independence growth (AIG)

AIG was determined in 6 well plates containing 2 layers of low melting point agarose in MEBM (Lonza) at 0.75% and 0.45% on top. 15000 cells/well were seeded and plates incubated for 5 weeks at 37°C in 5% CO2. Colonies were visualized using 0.01% cristal violet and counted. Prior staining colonies were isolated and put in culture to generate the Soft Agar (SA) clones.

### Tumorigenicity

1.5 or 5×10^6^ cells resuspended in a 1:1 Matrigel/PBS solution were injected subcutaneously in Swiss nude or SCID beige mice (Harlan/Envigo, Garnat France). Tumor growth was monitored for 5 months before animals were sacrificed. In vivo experiments were systematically reviewed and approved by an internal animal ethics committee and the University of Montpellier animal ethics committee.

### DNA and RNA extraction

DNA and RNA were isolated using the QIAmp DNA Mini kit and Rneasy Mini Kit (Qiagen S.A. France, Courtaboeuf, France). Each DNA sample was quantified by nanospectrophotometry (NanoView, GE Healthcare, Orsay, France) and qualified by 0.8% agarose electrophoresis. Qualification of mRNA was performed using a Bioanalyser (Agilent, Santa Clara, CA, USA).

### Array-CGH and mRNA expression profiling

Array-CGH was done using HG18 CGH 385K Whole Genome v2.0 array (Roche NimbleGen, Madison, WI, USA). DNA from a pool of 20 normal females was used as reference. For hybridization, 1 mg of genomic DNA and reference DNA were labeled using NimbleGen Dual-Color DNA Labeling Kit (Roche Diagnostics, Meylan, France). Labeling products were precipitated with isopropanol and resuspended in water. Test (Cy3) and reference (Cy5) samples were combined in 40 ml of NimbleGen Hybridization buffer. Hybridization was performed in a NimbleGen Hybridization system 4 for 48 h at 42 C with agitation mode B and washed using NimbleGen Wash Buffer kit according to manufacturer’s instructions. Arrays were scanned at 5 mm resolution using the GenePix4000B scanner (Axon Instruments, Molecular Devices Corp., Sunnyvale, CA). Data were extracted from scanned images using NimbleScan 2.5 extraction software (Roche NimbleGen, Madison, WI, USA), which allows automated grid alignment, extraction, normalization, and export of data files. Normalized files were used as input for the Nexus 6.1 Software (Biodiscovery, El Segundo, CA, USA). Analysis settings for data segmentation and calling were the following: significant threshold for FASTST2 Segmentation algorithm: 1.0E-7, Max Continuous Probe Spacing: 1000, Min number of probes per segment: 10, high level gain: 0.485, gain: 0.17, loss: 0.2, homozygous copy loss: 0.485. Hierarchical clustering was done using Nexus 6.1 using average linkage setting. Interval files from each of the 18 samples were exported and converted to a BedGraph format and merged (using the Merge BedGraph files tool (v 0.1.1, http://galaxy.sb-roscoff.fr) in order to define common interval between samples. Intervals smaller than 2 MB were removed resulting in a set of 268 intervals with CNA changes in at least one sample. This file was used as input of the MoCluster algorithm.

### miR expression profiling

Biotinylated cRNA were prepared according to the Affymetrix IVT Express protocol from 100 or 200 ng total RNA and hybridization was done as follows. CRNA were fragmented, 12 mg hybridized for 16 h at 45 C, washed and stained in the Affymetrix Fluidics Station 450 with Hybridization Wash & Stain kit. GeneChips were scanned using the Affymetrix GeneChip Scanner 3000 7G. Raw feature data were normalized using Robust Multi-array Average (RMA) method (R package affy). All subsequent analyses were performed on normalized datasets. To determine genes differentially expressed between sublines we used sam_multiclass command with one class for two biological duplicates (samr R package median). Lines containing identical GeneSymbol were collapsed with a max function. This file containing expression values for 18 samples and 2063 genes was used as input in MoCluster.

### DNA methylation profiling

RRBS libraries were prepared as previously described [47]. Genomic DNA was digested for 5 h with MspI (Thermo Scientific) followed by end-repair, A- tailing (with Klenow fragment, Thermo Scientific) and ligation to paired-end methylated adapters (with T4 DNA ligase, Thermo Scientific) in Tango 1X buffer. We purified fragments in the range 150 to 400 bp by electrophoresis on a 3% (w/v) agarose 0.5X TBE gel with the MinElute gel extraction kit (Qiagen), and performed two rounds of bisulfite conversion with the EpiTect kit (Qiagen). RRBS libraries were generated with PfUTurbo Cx hotstart DNA polymerase (Agilent) and indexed PE Illumina primers using the following PCR conditions: 95°C for 2 minutes, 12 to 15 cycles (95°C for 30 s, 65°C for 30 s, 72°C for 45 s), 72°C for 7 minutes. The libraries were purified with AMPure magnetic beads (Beckman Coulter) and sequenced (2 × 75 bp) on an Illumina HiSeq2000 by Integragen SA (Evry, France) to generate between 20 and 30 million pairs of reads per sample. The processing of reads was performed as described (Auclair et al, 2014). We aligned reads to the human genome (hg19) with BSMAP and only retained the CpGs sequenced at least 8X.

Determination of Differentially methylated regions (DMRs) between sublines was restricted to CGI comprising at least 5 contiguous CpGs, filtered for methylation differences of at least 0.2 between any of the samples. The frequency of the methylation levels was calculated and displayed as histogram in 40 classes to evaluate whole genomic variation of the DNA methylation (figure 3A). Then we used sam_multiclass function (samr R package median FDR= 0.0449). The result was a file of 892 DNA segments that was used as an input for MoCluster. Accordingly, we explored global DNA methylation variation genome-wide. In the expression correlation analysis we selected DMRs close to TSS (+/- 1000 bp to TSS).

### MoGSA

To integrate the omics data of different origins (CNA, mIR and Mrna and DNA Methylation), we used the MoGSA package (Meng, 2017a,b) to identify Joint Patterns Across Multiple Omics Data Sets. Mogsa have proved to be particularly efficient in term of computational time and compared favorably to icluster (Meng et al., 2016). We used consensus PCA from the MoGSA R package and displayed the results from the first and second principal component either for each omic dataset (CGH, DNA, mRNA or miR) or all together. The features with highest coefficient in the definition of the first axis of the PCA were selected and submitted to unsupervised Ward clustering and presented as heatmaps. Features varied according to the analysis and were as follows; Coefficient higher or equal to 0.07 in Fig 2C, coefficient higher or equal to 0.05 in Fig 3C, coefficient higher or equal to 0.07 in Fig 4b, coefficient higher or equal to 0.06 in Fig 4D.

### Immunofluorescence

Cells were seeded onto glass slides and grown to reach 50-60% confluence. Prior immunofluorescence cells were fixed with either 2% paraformaldéhyde or with ice cold methanol and permeabilized with 1% PBS-Triton and rinsed with 2% serum-PBS before incubation with the primary antibody, being rinsed in 2% serum-PBS and stained with DAPI and incubated with the secondary antibody. γH2AX/53BP1 foci were counted in triplicate on 400 nuclei. Antibodies are listed in the S Appendix.

### High throughput data analysis

Detailed methods are presented in the Supplementary Methods. Raw array and RRBS data can be accessed at GSE114849 (see S Appendix).

## Supporting information

**S1 Fig: Gradual immortalization and transformation of the shp53 HMEC sublines**. A, b-galactosidase staining; B, C: HTERT mRNA expression and telomerase enzymatic activity, the shp53 HTERT cells are presented as a positive control of TERT expression; D: anchorage independent growth in soft agar, number of positive experiments out of number of attempts, pictures of the corresponding soft agar petri dish and blow up of foci formed by the respective sublines.

**S2 Fig: Attenuation of p53 after shRNA transduction and overexpression of oncogenes in oncogene transduced sublines**. A: attenuation of p53 was ascertained by challenging primary R2 and R2-shp53 cells with Bleomycin for 6 hours, accumulation of p53 and induction of p21 were used as read outs for p53 functionality. B: QPCR verification of WNT1, CCNE1 and RAS mRNA expression in R2-shp53-WNT1, R2-shp53-CCNE1 and R2-shp53-RAS respectively. C: protein expression levels by western blotting. E: early; L: late.

**S3 Fig: Spontaneous DNA damage in shp53 sublines**. A: gammaH2Ax and 53BP1 staining patterns, note the pannuclear H2Ax staining in shp53 HMECs, whereas shp53-WNT1, shp53-CCNE1 and shp53-RAS show predominantly H2Ax foci. Hydroxyurea treatment was used as a control of H2Ax staining as a condition of severe replication stress. It is of note that primary R2 HMEC presented a sizeable number of foci positive cells, which can be attributed to telomere attrition in these cells. Bleomycine treatment was used as a control of double strand breaks. B: fraction of cells showing H2Ax pan-nuclear staining or more than 4 foci. C: Statistics of Neutral CometAssay tail moment measurement for 3 independent experiments in shP53 and shp53-oncogene sublines (top table) and Annova multiple comparison test (bottom table). shP53-CCNE1 and shP53-WNT1 sublines present a significantly higher number of double strand breaks than shP53-RAS. D: Tail moment measurements box plot.

**S4 Fig: CNA plots of HMEC models**. A: CNA at different steps of transformation. CNAs are represented for each chromosome, red for losses, blue for gains. The hight of the bars indicates the probability of occurrence. B: cumulated CNA plots of HMEC models. CNAs are represented for each chromosome, red for losses, blue for gains. The hight of the bars indicates the amplitude of the copy number change.

**S5 Fig: pathways and regulation networks principally affected in the different HMEC sublines.**

**S6 Fig: phenotypic characteristics of shp53 HMEC sublines**. Cells were stained by immunofluorescence (IF) for the expression of ECAD (E-Cadherin, green) which is a marker of epithelial cells, VIM (Vimentin, red) marker of mesenchymal cells, CK8 (Cytokeratin 8) marker of luminal breast epithelial cells, CK5 (Cytokeratin 5) marker of basal breast epithelial cells. Differential expression patterns can be observed according to the genetic elements expressed and the stage of the culture. Normal HMEC and Early shp53 co-express ECAD and VIM and are mosaiec for CK5 and CK8 expression. In shp53 late HMEC tended to lose ECAD expression and become mesenchymal, but kept a mosaic CK5/CK8 pattern. In shp53-WNT1 and shp53-CCNE1 ECAD and VIM were co-expressed in all cells indicating the conservation of an epithelial phenotype. Interestingly, whereas shp53-WNT1 expressed only CK5 and no CK8, shp53-CCNE1 preserved the original mosaic phenotype of the shp53 HMECs. Expression of RAS-v12 produced drastic changes as illustrated by the concomitant loss of expression of ECAD and both CK5 and CK8.

**S7 Fig: Genes modified by CNA or differential methylation in R2^shp53-RAS^ and R2^shp53-CCNE1^/R2^shp53-WNT1^ show little overlap**. Of note MTUS1 is strongly underexpressed in RAS (−2,95;about 1/10 x) whereas it is overexpressed in CW (0,87, about 1,8 x). PLAG1 is strongly overexpressed in RAS (3,25; about 9,5 X) and moderatly in CW (0,84 about 1,8 x) similarly to THRA (RAS change =1,89 about 3,7x; CW change =0,89 about 1,85x). ZNF7 expression change is equivalent in both clusters.

**S8 Fig: HMEC models show variable association to either the Basal A or Basal B subtype.** A: Classification was done using the BasalA/B classification the gene centroid list http://rock.icr.ac.uk/collaborations/Mackay/centroid.correlations.Eset/ExpressionSetNearestCentroidCorrelations.pdf) (Neve et al., 2006) and spearman correlation. R2 (Ctrl here) and R2shp53 showed co-correlation with Basal A and Basal B. R2shp53-RAS models showed strongly positive correlation to Basal B and negative correlation to Basal A, whereas R2shp53-CCNE1 and R2shp53-WNT1 showed stronger correlation to Basal A. B: radar plot of the mammary differentiation scores as defined by E. Lim et al (2009) of our HMEC models. The different HMEC sublines presented a high luminal progenitor and luminal mature score with little variation among samples. Of note the elevated stromal score of shp53-RAS HMECs and the globally marginal MaSC score.

**S9 Fig:** Principal pathways activated in claudin-low TNBC

**S1A Table: genes with copy number dependent expression changes in the shp53-ras cluster vs. The shp53 and R2 cluster.** Genes were selected after a Spearman correlation test on the complete dataset and 90 genes from this comparison corresponded to genes with log2 scale changes (copy number change <- 0.15 or > 0.15; expression change <-0.5 or expression change >0.5; uncorrected t-test for expression data <0.05)

**S1B Table: genes with copy number dependent expression changes in the shp53-ccne/wnt cluster vs. The shp53 and R2 cluster.** Genes were selected after a Spearman correlation test on the complete dataset and 156 genes from this comparison corresponded to genes with log2 scale changes (copy number change <- 0.15 or > 0.15; expression change <-0.5 or expression change >0.5; uncorrected t-test for expression data <0.05).

## Acknowledgments

Authors wish to thank Anne Morel and Laurent Le Cam for providing retroviral constructs used in this project and Alexandre Djiane for critical reading. The expert input of the personnel of the microarray Center of the Genomics Research Unit Luxembourg Institute of Health is gratefully acknowledged.

## Authors contribution

CF, AS, AAK: cell and molecular biology, retroviral transductions, biochemistry. BO: array-CGH and data analysis. ES, JC: biomathematical analyses of gene expression data. EC, AB, MD, MW: RBBS libraries and DNA methylation data production. WJ, CS: helped design and edited the manuscript. SdM: designed and performed bioinformatics and biomathematical analyses, edited the manuscript. CT: Study design and coordination, fund raising, manuscript preparation and writing.

